# Stochasticity in Dietary Restriction-Mediated Lifespan Outcomes in *Drosophila*

**DOI:** 10.1101/2024.09.06.611756

**Authors:** Olivia L. Mosley, Joel A. Villa, Advaitha Kamalakkannan, Eliyashaib James, Jessica M. Hoffman, Yang Lyu

## Abstract

Dietary restriction (DR) is widely considered to be one of the most potent approaches to extend healthy lifespan across various species, yet it has become increasingly apparent that DR-mediated longevity is influenced by biological and non-biological factors. We propose that current priorities in the field should include understanding the relative contributions of these factors to elucidate the mechanisms underlying the beneficial effects of DR. Our work conducted in two laboratories, represents an attempt to unify DR protocols in *Drosophila* and to investigate the stochastic effects of DR. Across 64 pairs of survival data (DR/ad libitum, or AL), we find that DR does not universally extend lifespan. Specifically, we observed that DR conferred a significant lifespan extension in only 26.7% (17/64) of pairs. Our pooled data show that the overall lifespan difference between DR and AL groups is statistically significant, but the median lifespan increase under DR (7.1%) is small. The effects of DR were overshadowed by stochastic factors and genotype. Future research efforts directed toward gaining a comprehensive understanding of DR-dependent mechanisms should focus on unraveling the interactions between genetic and environmental factors. This is essential for developing personalized healthspan-extending interventions and optimizing dietary recommendations for individual genetic profiles.

## Introduction

Over the past century, the benefits of caloric or dietary restriction (CR or DR) have been extensively studied across organisms ^1,2^. The concept that reducing food intake without causing malnutrition may promote longevity and health is widely appreciated and generally supported by observations across various species. This field was anchored in early studies of McCay et al. ^3^, who reported that rats on a calorically restricted diet were longer lived than those fed ad libitum (AL). Since then, the effects of CR/DR have been demonstrated to extend to multiple species including yeast ^4^, invertebrates ^5,6^, other mammals ^7^, and perhaps even humans ^8^. Remarkably, the underlying biology of CR/DR reveals a complex and conserved molecular machinery, that includes pathways that play a crucial roles in nutrient sensing and DR-mediated outcomes such as the target of rapamycin ^9^ and AMPK-activated protein kinase pathways ^10^ (reviewed in ^2^).

While many would argue that DR is the most robust method to extend healthy lifespan known thus far, the complex nature of lifespan modulation under DR has become increasingly evident as genetic factors and other variables have been suggested to play significant roles ^11^. For instance, grand-offspring of wild-caught mice had no increase in longevity under DR ^12^, and less than 50% of 41 recombinant inbred mouse strains subjected to DR exhibited an increase in lifespan ^13^. More recently, Wilson et al. utilized 161 isogenic strains from naturally derived inbred lines of *Drosophila melanogaster*, finding that 29% of these strains did not exhibit DR-induced lifespan extension ^14^. These findings underscore the need to further investigate and explore influential variables, including but not limited to genetic background, to enhance our understanding of the relationship between DR and longevity control.

In addition to genetic factors associated with response to DR, stochastic events are increasingly recognized as significant contributors to the diversity of aging phenotypes ^15-17^. For example, *C. elegans* from an N2 isogenic reference population show varied rates of aging as they approach later life stages ^18^, and the *Caenorhabditis* Interventions Testing Program (CITP) has found significant stochastic variation in lifespan across and within laboratories ^19^. In flies, stochastic variation has been observed in response to mating status across genetically distinct population ^20^. Furthermore, recent studies have identified intrinsic noise and variations at the cellular level in aging biomarkers ^21,22^. Overall, the inclusion and rigorous analysis of stochastic factors in DR studies are critical and currently underexplored, potentially biasing results of DR experiments.

Invertebrate models such as *Drosophila* and *C. elegans* have been instrumental in elucidating key factors that contribute to the longevity benefits of DR. These models have primarily explored DR by modulating nutritional concentrations in the food media ^23^, not necessarily restricting calories. Therefore, the AL state is better described as a high nutrient state, as both the DR and AL groups have continuous access to food. In *Drosophila*, restrictions of either yeast (a major protein source for flies) or individual amino acids have been extensively used to study DR mechanisms e.g. ^6,9,24,25^, though these studies have sparked some recent controversies (see recent updates from ^26^). Notably, the effects of dietary restriction are more consistent when a restricted diet is compared to a nutrient rich diet, rather than to a standard husbandry diet ^23,27^, though within *Drosophila* there is actually no “standard diet” used consistently across laboratories. This practice in the field presents significant challenges in attributing longevity effects solely to DR, as it has been shown that an enriched diet can lead to desiccation causing increased mortality ^27^, and overnutrition with a nutrient rich media may lead to obese phenotypes which predictably exhibit a shortened lifespan.

We suggest that the subtleties between a restricted diet and a “standard” diet may present challenges in reproducibility due to stochastic variations, and that DR effects may only be biologically relevant when compared to high nutrient, enriched diets. To assess and quantify these variations, we replicated DR experiments that involve multiple cohorts and distinct dietary paradigms, in two geographically distinct laboratories. We find that while genotype emerges as the most significant predictor of lifespan, we recorded considerable variation among cohorts with respect to DR effects, some of which can be attributed to stochastic variation. We conclude that rigorous understanding of CR/DR outcomes must strongly take genetics and stochastic factors, as well as diet details, into account.

## Methods

### *Drosophila* husbandry

Mated male and female flies from four common laboratory strains of *Drosophila melanogaster* were used in each cohort: *w*^1118^, Oregon-R (OR), *w*^Dahomey^, and Canton-S (CS). As an additional control for any potential genetic drift or variations between stocks, the Hoffman lab gifted OR and w^1118^ strains and received the *w*^Dahomey^ and Canton-S strains from the Lyu lab, so the strains used across labs were genetically identical. After exchange, all new fly strains were acclimated to the laboratory for a period of 6-8 weeks prior to use in experiments. Lab stocks were maintained at 25°C at 65-85% humidity and a diurnal, 12-12 light/dark schedule. All fly stocks were maintained on a cornmeal-based (CT) diet (Table 1).

**Table 1.** Ingredients of each of four diets used in the study. Each amount is measured in 1L of water. The nutrient composition is estimated using *Drosophila* Dietary Composition Calculator: https://brodericklab.com/DDCC.php.

| Ingredient | Experimental Protocol 1 |  | Experimental Protocol 2 |  |
| --- | --- | --- | --- | --- |
|  | DR (CT) | AL (SY10) | DR (SY5) | AL (SY15) |
| Agar | 1-2% | 1-2% | 1-2% | 1-2% |
| Propionic acid (mL) | 5 | 5 | 5 | 5 |
| Yeast (g) | 25 | 100 | 50 | 150 |
| Sucrose (g) | 55 | 100 | 100 | 100 |
| Dextrose (g) | 30 | 0 | 0 | 0 |
| Cornmeal (g) | 60 | 0 | 0 | 0 |
| Total calories (cal) | 628.85 | 775.70 | 582.20 | 969.20 |
| Proteins (g) | 17.72 | 53.03 | 26.53 | 79.53 |
| Fat (g) | 1.44 | 0.30 | 0.30 | 0.30 |
| Carbohydrates (g) | 150.43 | 151.00 | 129.50 | 172.50 |

### DR lifespan protocols

Both labs collected time-synchronized eggs for the lifespan assays. In the Hoffman lab, each genotype was placed on fresh CT food, and flies mated and laid eggs for 48-72 hours. After expanding each stock, all adult flies were cleared from the vials and the time-synchronized eggs developed. The Lyu lab used an egg-collecting chamber and grape juice-agar media to gather embryos deposited within a 48-hours period ^28^. For both labs, after 10 days, the new adult flies were transferred onto SY10 (Cohort 1, 3, and 4) or CT (Cohort 2) food and allowed to mate for 48 hours before sexing under light CO_2_ anesthesia. The difference in the mating diet introduces variation in early life dietary exposures. The collection process took place over the course of 2-3 days until 300 flies were collected for each genotype and sex with each vial containing 25 flies. The collected flies were randomized onto either a dietary restriction (DR) or ad libitum (AL) media (Table 1). We must note that while we are using the term ad libitum for the higher nutrient treatment due to the ubiquitous use of the term in the aging field, in *Drosophila*, and other invertebrates, this is not at true AL treatment, as all groups have access to their diet 24/7. We varied the diets and mating food in individual cohorts such that cohorts 1-3 used CT/SY10, while cohort 4 used SY5/SY15 as the DR/AL dietary paradigms, respectively. Flies were transferred to fresh media three times a week with deaths recorded at each transfer using D-Life ^28^ and Excel.

### Climbing and body mass assays

At approximately 30 days of age, flies from each group were run through a climbing assay. Briefly, flies were tapped to the bottom of an empty vial and allowed to climb for 10 seconds. At 10 seconds, the number of flies that had climbed at least 5 cm was recorded. Data was collected from cohorts 1-3 in the Hoffman lab and analyzed with all results combined.

To determine if flies on low yeast diets were calorically restricted, we weighed flies on each diet to determine if the DR flies weighed less than AL flies. Flies were placed on either a S10Y5 or S10Y15 diet for 30 days prior to weighing. After 30 days, flies were anesthetized on ice, transferred to a 2mL centrifuge tube in groups of 5-10 and weighed on a microanalytic balance. Weights were calculated by subtracting the average empty-tube weight per group from the measured weight per sample and adjusting for the number of flies per sample. Both climbing ability and body mass assays were only conducted in the Hoffman lab, as we were looking at general health effects, not reproducibility.

### Statistical analyses

All statistical analyses were completed in program R. Overall, comparisons across labs and variables of interest were determined with Cox proportional hazard models using the “survival” package ^29,30^. Comparisons between individual DR pairs within a lab/genotype/sex/cohort were made with log rank tests. Kaplan-Meier curves were plotted for visualization of the data. Spearman rank correlations were calculated to look at correlations of median longevities across laboratories. Due to the large number of log-rank tests for individual comparisons, we applied a Bonferroni correction with significance set as p<0.00078. Differences in healthspan measures (climbing ability and body mass) were calculated using an ANOVA looking at the effects of sex, genotype, and dietary treatment.

We performed Cox regression and model fitting using in-house R script to determine the amount of variance explained by each variable analyzed. We used the coxph function from the survival package ^29,30^ to fit both full and reduced models. The full model included the covariates lab, sex, cohort, genotype, and diet while the reduced models excluded one covariate at a time to evaluate their individual contributions. The proportional hazards assumption for the Cox regression models was tested using the cox.zph function from the survival package. We estimated the Cox-Snell R^2 31^ for both full and reduced models. The likelihood of each model was computed using the logLik function from the stats package. The contribution of each covariate was estimated using a likelihood-based measure, derived from the differences of log-likelihoods of the full and reduced models:

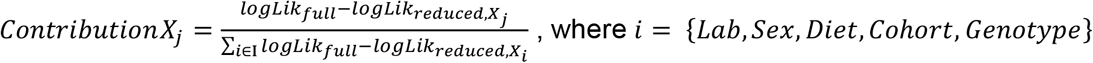

## Results

### Lab reproducibility

To minimize inter-laboratory variability and enhance reproducibility, we utilized the same DR protocols, applied identical experimental procedures, and ordered supplies simultaneously from the same vendors. Detailed approaches are described in the methods section. Our final dataset consisted of 15,935 flies across 64 pairs of DR/AL survival data (128 longevity curves). All raw data can be found in Supplementary Table 1. We used two DR protocols: Protocol 1 utilized the commonly used CT food as the restricted diet and SY10, 10% (w/v) sucrose:yeast as the AL diet, while Protocol 2 controlled for all other ingredients, varying only the concentration of yeast to further test the effects of protein restriction (see Table 1 for detailed ingredients). We ran Protocol 1 three times independently in each lab. We combined data generated from two protocols to estimate overall reproducibility and stochasticity.

Overall, we found reasonable reproducibility in lifespan data from the two labs (Figure 1a). We did find a significant difference in longevity between labs (log-rank p=0.001); however the differences in median lifespan are minimal: 53.7 days (95%CI 53-54.1 days) for the Hoffman Lab and 53.1 days (95%CI 52-54.2 days) for the Lyu Lab, a difference of ∼1% and driven by our large sample size (n = 8,475 for the Hoffman Lab and 7,460 for the Lyu Lab). Across cohorts, there was significant correlation of mean longevities between the labs (Figure 2, Spearman rho=0.55, p=3.6×10^−6^). Together, these results indicate that when applying the same protocols and procedures, laboratories or geographic locations are not major factors influencing lifespan results.

**Figure 1.**
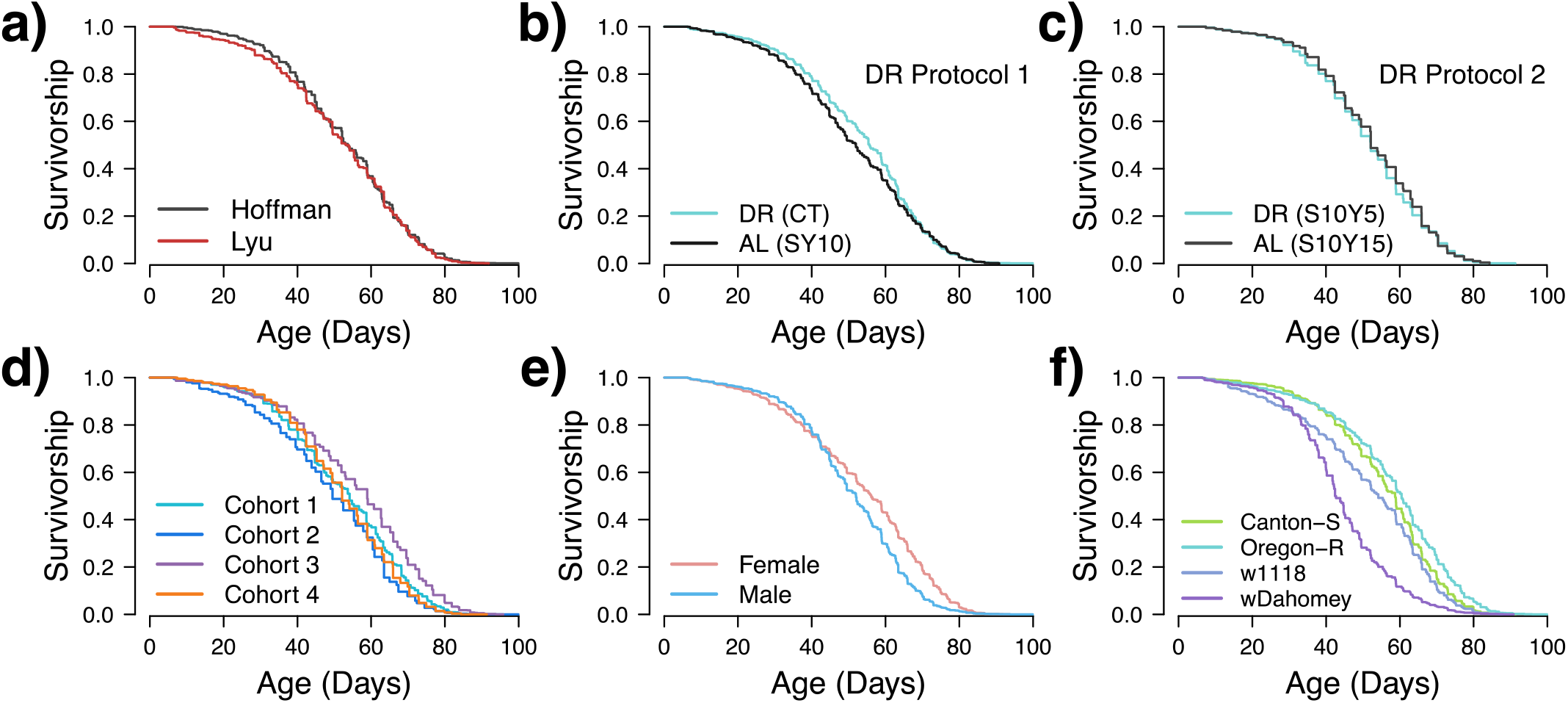
Kaplan–Meier survival analysis of *Drosophila melanogaster*. The survival curves represent the proportion of survivors over time (in days) during the adult stage. Each panel shows the survival curves for a specific factor, illustrating the effects of lab (a), dietary restriction protocol 1 (b), dietary restriction protocol 2 (c), cohorts (d), sex (e), and genotype (f) on the lifespan of the flies. These panels collectively demonstrate how different factors impact the lifespan of the flies.

**Figure 2.**
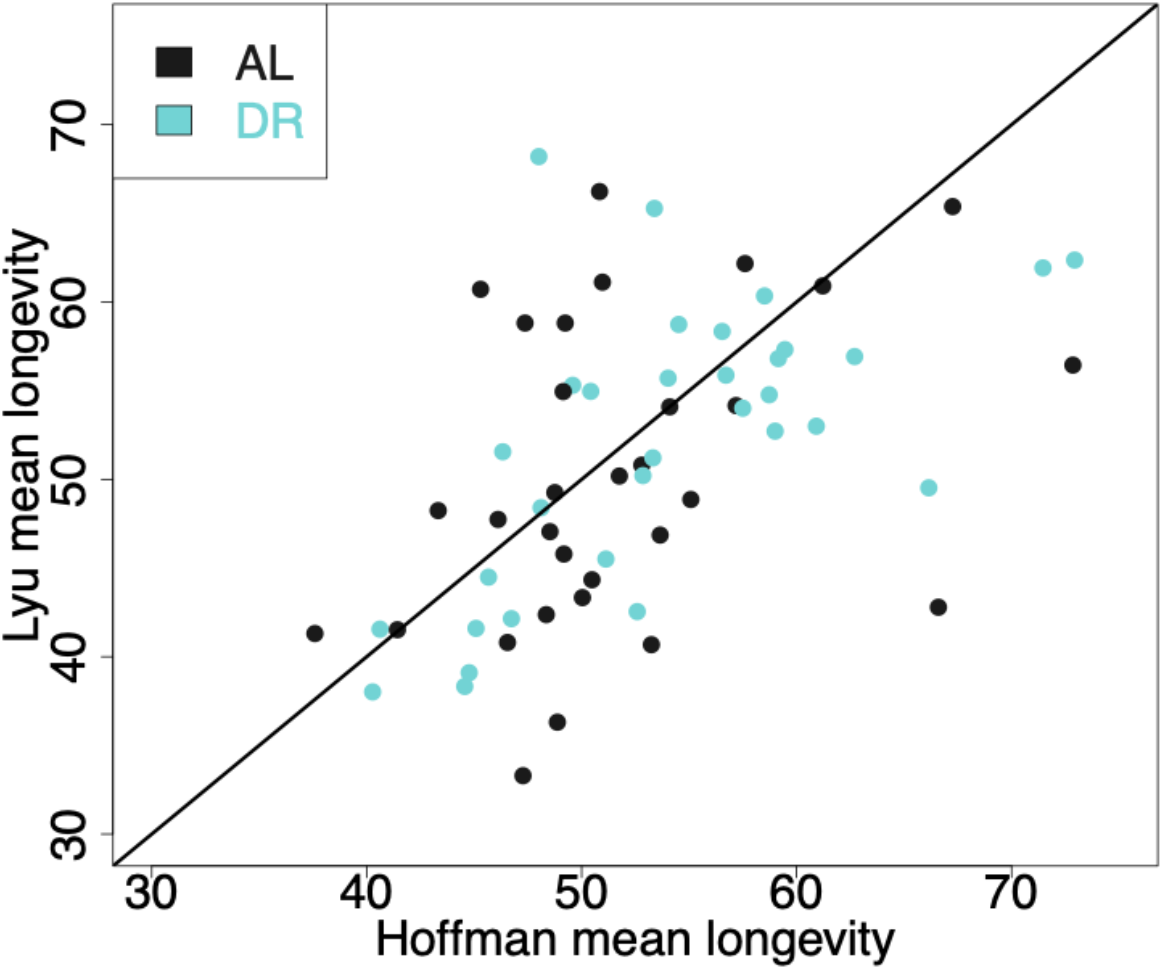
Correlation plot of each set of cohort pairs between the two labs. Each point represents the mean longevity for the Hoffman lab (x-axis) or Lyu lab (y-axis) for each individual treatment, sex, genotype, lab, cohort replicate (n=64). Black line is the line of symmetry.

### Biological factors and stochasticity together influence lifespan

To understand how each factor influences lifespan, we used Cox regression to estimate the predictive power of each factor in the model fitting (see Methods for detailed calculation and Discussion for limitations). Specifically, we calculated the Cox-Snell R^2^ for both the full model and the reduced models to examine the power of each covariate. We found that, compared to the full model, only removing the covariates genotype or cohort resulted in a moderate reduction in Cox-Snell R^2^ (Supplementary Table 2), indicating that genotype and cohort are the main factors influencing lifespan in our dataset.

We further estimated the proportion of variance each factor explains using a likelihood-based method, summarized in Table 2. The variability among cohorts (16.35% of total variance) indicates the presence of stochasticity, which explains the results even better than sex (14.59% of total variance), a well-known factor that determines lifespan ^32^. Genotype was the major determinant of variation in our dataset, accounting for 67.97% of the total variance. The variability between labs (3.33% of total variance) is small, consistent with our previous observation. Most surprisingly, dietary treatment, the main focus of this study, accounts for only 0.76% of the total variance. To visualize the differences, we present the average lifespan grouped by each factor in Figure 1, highlighting the negligible differences between labs (Fig. 1a) and dietary conditions (Figs. 1b and 1c), moderate differences in sex (Fig. 1e) and cohort (Fig. 1d), and remarkable differences in genotype (Fig. 1f).

**Table 2.**
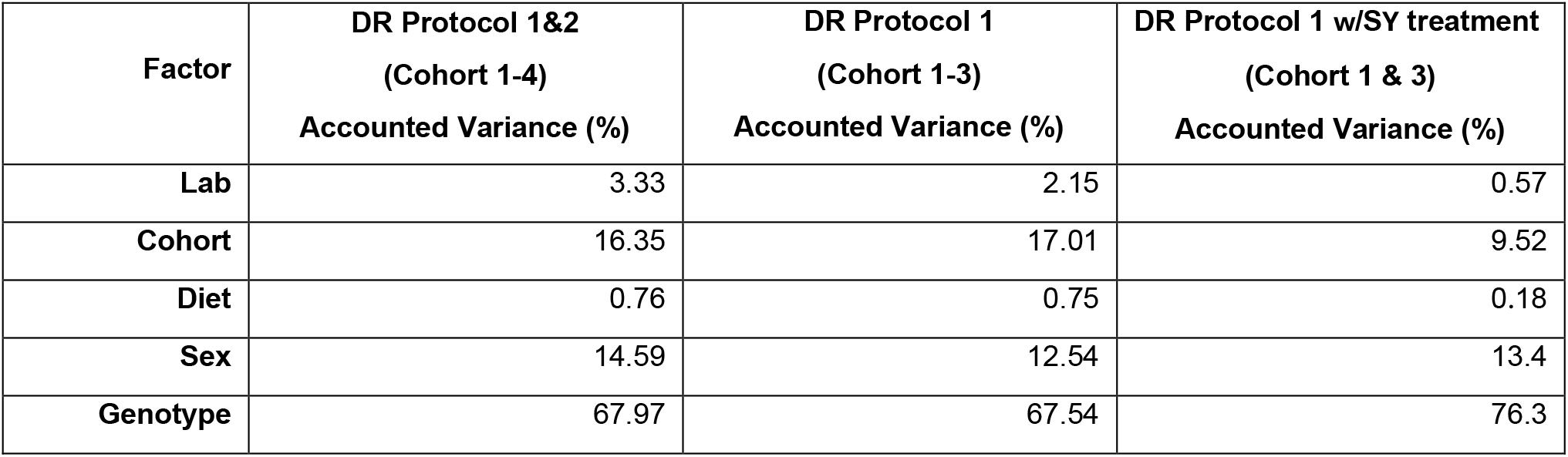
We estimated the accounted variance percentages for different factors across two dietary restriction (DR) protocols involving different cohorts, using a likelihood-based method.

We consider the possibility that different DR protocols might contribute to the stochasticity, even though this is not suggested by data in Figure 1b and 1c. To rule out impact of different DR protocols, we estimated the proportion of variance explained using DR Protocol 1 (Cohorts 1-3), which shows a similar result to the entire dataset, indicating that stochasticity may account for 17.01% of the total variance. We also suspect that different food flies mated on before the lifespan assay (Cohort 1 and 3 versus 2, see Methods for details) may add to the stochasticity. To test this, we estimated the proportion of variance explained with only Cohorts 1 and 3, where the food flies mated on are the same (SY10). Indeed, we observed a decrease in the proportion of contribution by cohort (9.52%), but this number is still much larger than the proportion contributed by lab (0.57%) and diet (0.18%). In summary, our analyses indicate that genetic, sex, and stochastic factors are the predominant determinants of lifespan, with lab and dietary restriction regimen accounting for very little impact on longevity.

### DR does not universally extend lifespan

One of the primary objectives of our experiment was to assess the reproducibility and stochastic nature of the longevity effects observed with dietary restriction. Combining two protocols, we found that DR flies were significantly longer-lived than those on high nutrient diets (Log-rank p=4.7×10^−7^), but the difference in median lifespan (7.1%) is rather small. Given the large stochastic effects in our dataset, we asked if the DR effects are reproducible across different replicates. Out of the 64 pairs of DR/AL comparisons, we observed a significant lifespan effect of dietary restriction in only 17 out of 64 pairs (26.7%, log-rank test, Table 3). Survivorship curves are shown in Figure S1. Unexpectedly, in five comparisons, the AL group exhibited significantly longer lifespans. Previous research has consistently indicated that *D. melanogaster* tend to live longer under dietary restriction (Grandison, Wong et al. 2009; McCracken, Adams et al. 2020). However, our findings can be extrapolated to suggest that these effects are at least partially attributable to the toxicity of the enriched diet (see Discussion). The effects of dietary restriction *per se* appear to be minimal and sporadic when compared to what would be considered a standard diet.

**Table 3.**
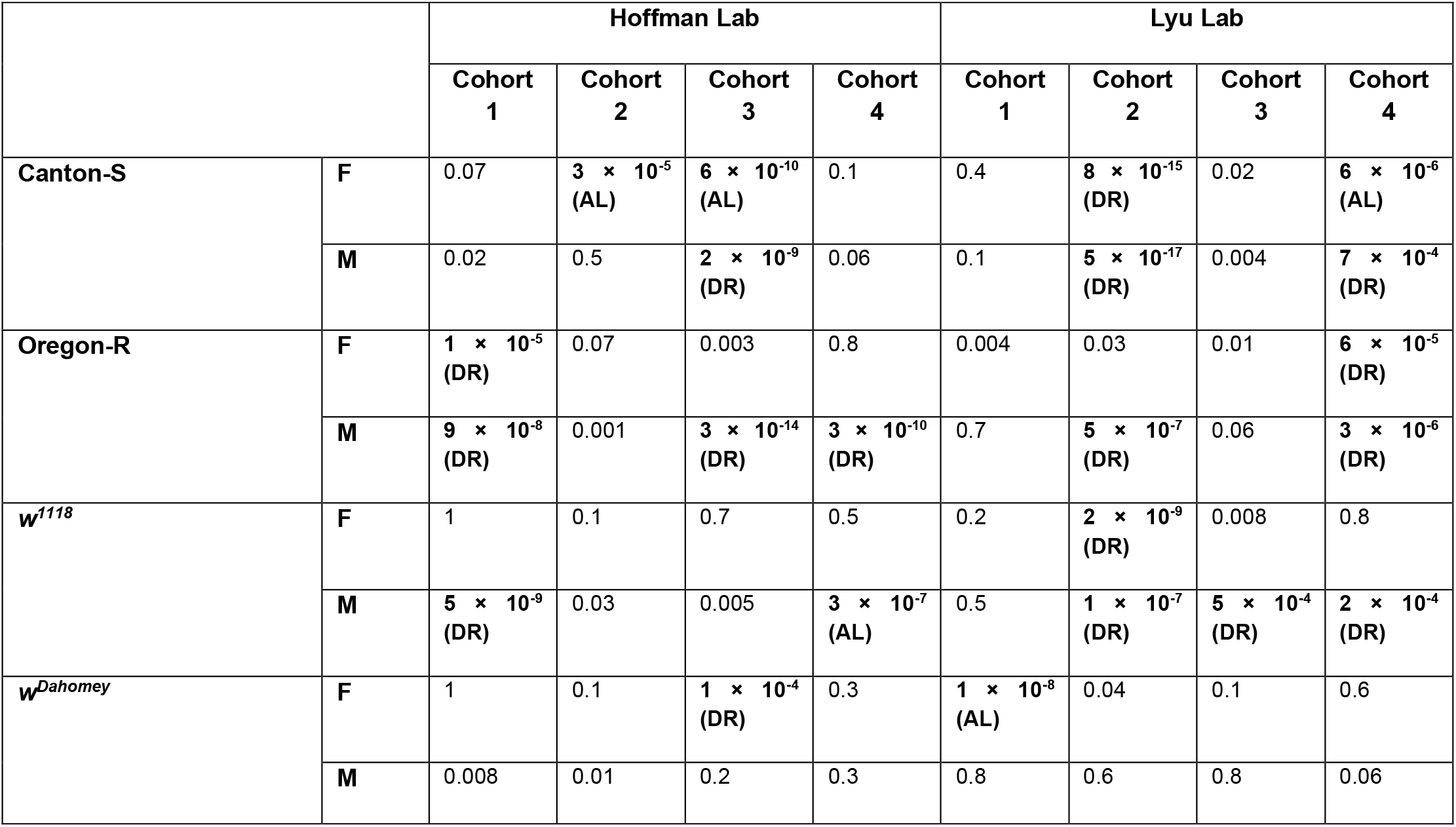
Log-rank test results and *P*-values for each AL/DR comparison pair. Bold text indicates *P*-values that pass the Bonferroni correction (*P* ≤ 0.00078). The longer-lived group is indicated in brackets for significant comparisons.

Within these analyses, certain genotypes were more likely to show an effect of DR, with *w*^Dahomey^ flies showing overall no real effect of DR, and Oregon-R flies showing lifespan extension under DR in almost 50% of the replicates with no increases in the AL groups (Table 3 and Figure 3a). Lastly, we also found significant sex-by-genotype effects, and in general males were more likely to respond to the different diets (Figure 3b), with the exception of *w*^Dahomey^, in which no male replicates had any significant differences between the DR and AL diets.

**Figure 3.**
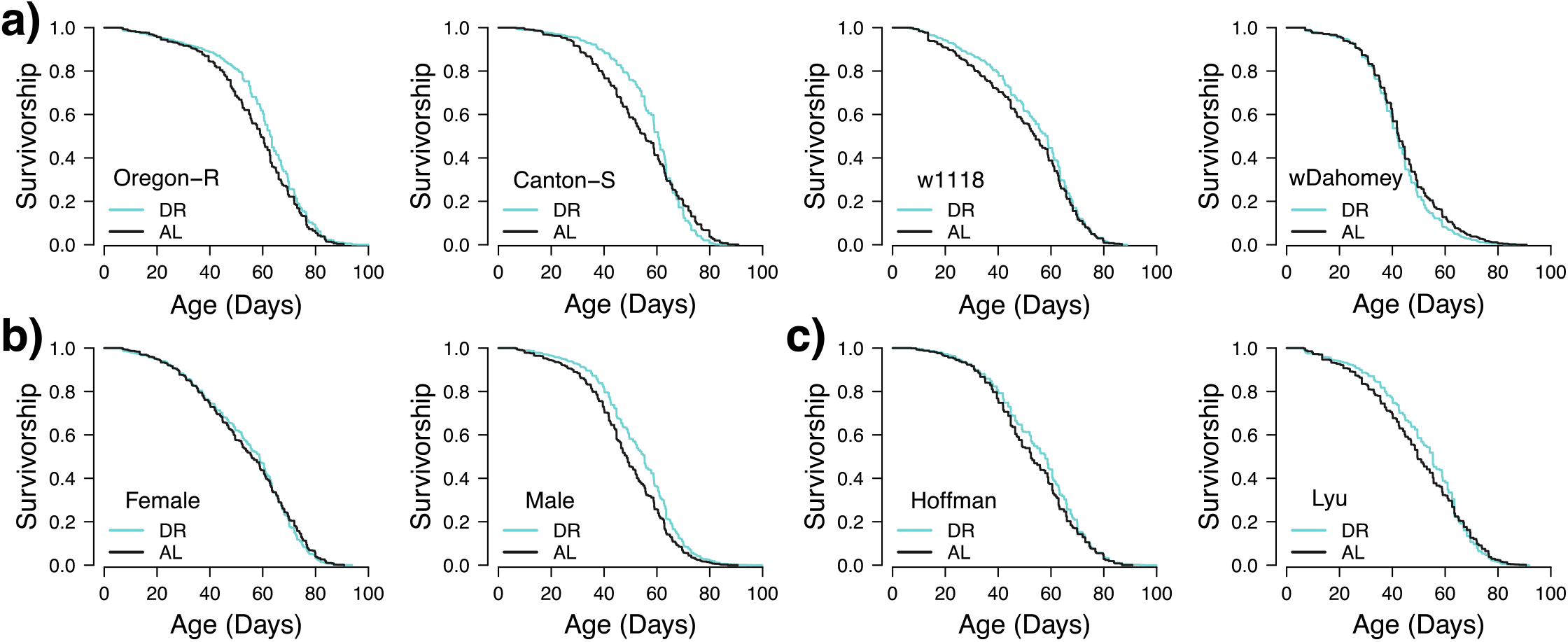
The interaction of DR with sex (a), genotype (b), and lab (c). Kaplan–Meier survival curves represent the proportion of survivors over time (in days) during the adult stage. Only cohorts 1-3 (DR protocol 1) were used for this analysis to control for the protocol.

We also observed variation in the two DR protocols, with Protocol 1 showing more pronounced DR effects (Figures 1b versus 1c). When examining the two labs separately, the Lyu lab found more DR effects using DR Protocol 2 (4 out 8 pairs have *P* ≤ 0.00078). In addition, looking at the difference between AL-DR median lifespans, we see a positive trend for similar differences between pairs, though the effect was not significant (Supplemental Figure 2, Spearman rho=0.313, p=0.081). Given the stochastic effects observed between different cohorts, it is difficult to determine whether these differences are due to variations between the labs or stochasticity.

Similar to our longevity results, we found no effects of DR treatment on climbing ability, a marker of healthspan, in middle aged flies (ANOVA p=0.54, Supplementary Figure 3), but similar to the longevity results, there were significant effects of genotype (ANOVA p=1.07×10^−7^) and sex (ANOVA p=0.004). As expected, our body mass analysis found males were smaller than females (ANOVA p=1.32×10^−7^, Supplemental Figure 4), but there was no effect of DR treatment (ANOVA p=0.23) nor genotype on overall body mass (ANOVA p=0.43). These combined data suggest there were no effects of our DR protocols on healthspan in the flies.

## Discussion

Although DR is widely acknowledged as one of the most effective pro-longevity interventions across various species, recent studies from several species indicate variable lifespan responses to restricted diets ^13,14,27^. These variations can be attributed to differences in dietary regimes, genetic backgrounds ^27^, laboratory conditions ^33^, and other stochastic effects. Our collaborative effort, involving the replication of identical sets of experiments between two labs and repeating the same experiments multiple times within each lab, provides a unique opportunity to focus on the stochastic effects of DR. While we observe variations in lifespan between labs, these are not necessarily greater than the variations seen within repeated experiments in the same lab. This finding suggests that with rigorous control of laboratory conditions, inter-laboratory variability can be minimized (less than 4% of the total variance in our dataset), allowing a clearer focus on biological and stochastic effects, and our study strongly suggests that stochastic effects are one of the primary variables influencing lifespan under DR (Table 2). This conclusion is consistent with major findings from the *Caenorhabditis Interventions Testing Program* (CITP) ^19,34,35^, supporting the notion that variability in longevity control might be a universal phenomenon. Therefore, while DR may be a robust method to increase lifespan, there is significant variation in the magnitude and directionality of response.

One of the key reasons we observed significant stochasticity in our results is perhaps the small average lifespan differences between the DR and AL conditions in most of the genotypes, even when the sample size is sufficiently large (Figure 1b and 1c, Table 3). The average lifespan response to varying protein (yeast) concentrations in the diet typically follows a bell-shaped curve across different genotypes ^27^. A major challenge in designing DR experiments is determining the optimal food formulation that maximizes lifespan under DR conditions, as well as identifying an appropriate standard diet for the high nutrient group. A common misinterpretation of DR effects arises when using an extra high-nutrient diet as the control, often referred to as the AL condition. In such cases, observed lifespan extensions under DR could be misleading, as they may reflect the harmful effects of a high-nutrient diet rather than true benefits of DR (see Discussion in Ref. 27). For example, recent studies suggest a large effect of DR on *Drosophila* lifespan ^36^, but the AL diet was 30% (w/v) Y and S, which is well outside of what is used in standard husbandry. When a standard diet is chosen properly (e.g. 1% compared 10% (w/v) S and Y, as shown in ^37^, the differences between the DR and AL groups tend to be subtle in most genotypes, as observed in our study and reported by others ^23,27,38^. Given this perspective, the lack of significant DR effects, though initially unexpected, becomes less surprising. This subtlety in DR response emphasizes the importance of carefully selecting control diets and highlights the inherent challenges in designing and interpreting DR studies.

Although we did not observe a remarkable lifespan extension with DR, the differences between the DR and AL groups were reasonably repeatable across our labs (Fig. 2). A previous report has analyzed the correlation between the lifespan differences (DR-AL) in their dataset ^33^ and those by a second study ^39^, reporting a correlation, although not statistically significant. This lack of significant correlation could be influenced by variations in fly husbandry and dietary regimes between the studies ^33^. Nevertheless, the delta in lifespan between the DR and AL conditions (ΔL [DR-AL]) seems relatively consistent, even if the differences are not always significant nor positive (Supplementary Fig. 2). This “rule” suggests that within each genotype, the lifespan response curve to dietary concentration ^37^ is relatively stable.

Our findings underscore the importance of controlled experimental conditions and highlight the inherent challenges in achieving significant lifespan extensions through DR in certain genotypes. However, it is worth noting that the Oregon-R genotype consistently exhibits a DR response in 12 out of 16 trials in our studies (P < 0.05), with none showing an increase in the AL group. This suggests that in specific genetic backgrounds, the response to DR may be more predictable and robust. Understanding the genetic bases underlying this robustness is critical for future mechanistic studies, and for translating DR interventions into practical applications in daily life. Interestingly, we found remarkably similar median and maximum lifespans within a genotype across laboratories suggesting strong genetic effects on strain longevity, but not necessarily on strain response to DR. This is similar to our previous work suggesting high genetic correlation across strains within and between labs ^20^. Together, both genotype (G) and the interaction between genotype and diet (G x E) seem to have more significant impact on longevity than diet alone (E). Thus, as has been becoming more and more evident in the aging field, studies of multiple genetic backgrounds are necessary to understand the species level effects of different interventions and environmental conditions.

It may be noteworthy that we found no effect of diet on climbing ability or weight across our treatments, although these data we collected in only one of our labs (Supplementary Fig. 3 and 4). This suggests that first, what we are considering to be AL/DR in flies is not an accurate representation, specifically the AL group, as the DR group did not have a small body mass than the AL group, as would be expected in mammalian CR studies, where CR mice are significantly smaller than those on AL diets ^3,12^. Potentially we need a new way to denote DR studies that refer to the high/low nutrients of the diet but not necessarily the caloric intake of individuals on the diet as is denoted by the name ‘ad libitum’. In addition, as we found minor effects of DR on lifespan, it is not particularly surprising that health was also not affected. This is in line with previous studies showing that the correlation between health- and lifespan also depends on the genetic background ^14,34^. Like our longevity results, we found effects of sex and genotype on the climbing ability (and weight) that completely overshadowed any DR effect. Combined, these results suggest again that DR may have minor effects in *Drosophila* when restricted animals are compared to a ‘standard’ diet, and genetic background effects drive most of the variation in organismal health in fruit flies.

### Caveats

#### DR Protocols

While our results hint toward some of the nuanced conditions that must be considered when interpreting dietary interventions and longevity response in *Drosophila*, our results are not without their limitations. Experimental diets using CT and SY10 foods were selected based on their common use as stock diets in *Drosophila* laboratory husbandry. The addition of cornmeal in the CT food may slow the mechanical ingestion and metabolism and have physiological impacts, though our minor longevity effects seen comparing CT and SY10 suggest these effects are most likely minor. In addition, our SY5 and SY15 diets did not show many DR effects in the Hoffman lab specifically, suggesting our lack of CT/SY10 effects are most likely not due to any intentional differences in the food media. Both labs experienced issues with food quality across the experimental cohorts leading to censoring of flies, usually related to overly wet/sticky food; however, these food issues were random and would have been equally applied to all groups minimizing their overall effects. Still, we cannot rule out a bias in our removal of individual flies from the analysis.

#### Modeling

The assumption of proportional hazards in the Cox Regression model was not met, as indicated by the p-values from the proportional hazards test being less than 0.05 for all covariates except Lab. Given our large sample size, it is challenging to completely avoid violations of this assumption, and even small deviations can look like a violation when they are not biologically meaningful. Perhaps not surprisingly given the rest of our results, the largest deviations from the proportional hazards assumptions were due to the genotype effects. In the future, adjusting the model to include time-dependent covariates may address these violations and improve the accuracy of our results.

### Conclusions

Combined, our results find inconsistent DR longevity effects across labs within *Drosophila melanogaster*. As fruit flies are common longevity and dietary intervention models, it is important to note that any observed longevity effects in other studies may be due to stochastic variation within and across labs. We would suggest future studies need to thoughtfully design experiments with appropriate AL diets, and in addition, future studies must carefully interpret data, especially those that apply to minor effects. This caution is also likely relevant to other invertebrate species. Moving forward, one of the priorities perhaps should be focused on mapping the genetic alleles that influence the degree of variation in DR-mediated lifespan changes, as genotype was the largest factor affecting both overall longevity and response to DR. Utilizing existing population genomic resources will be essential in identifying such genetic determinants. Our insights on diet and longevity relative to genetic make-up, food regimen, and stochastic factors, will be crucial for advancing effective approaches for personalized medicine and nutrition, allowing for more tailored and effective longevity interventions.

## Acknowledgements

The authors would like to thank members of the Lyu and Hoffman labs for help with fly husbandry. We would also like to thank Monica Driscoll for her valuable comments on the manuscript. This work was funded by R00AG059920 to JMH.

## Data availability

Our lifespan data is available in Supplementary Table 1.

## Author contributions

The paper had the following contributions by each author: conception and experimental design-JMH and YL; data collection-OLM, JV, AK, EJ; data analysis, figure creation, writing first draft-OLM, JMH, YL. All authors edited and approved the final version of the manuscript.

## Supplemental Figure legends

**Supplemental Figure 1. Kaplan-Meier curves of each of 64 pairs of AL/DR experiments**.

**Supplemental Figure 2. Difference of AL and DR median lifespan between labs**.

**Supplemental Figure 3. Climbing results for 30-day old flies from the Hoffman lab for females (A) and males (B)**. Each replicate consists of 18 vials of ∼20 flies each. Mean climbing values were taken on a per vial average. Cohorts 1-3 were combined for analysis. There were significant effects of sex and genotype with no difference between AL and DR treatments.

**Supplemental Figure 4. Body mass results for 30-day old flies on SY5 vs SY15 for females (A) and males (B)**. Each replicate consisted of ∼ 5 measurements of 5 flies each. There were no significant effects of treatment, suggesting that our flies were not calorically restriction on the DR treatment. Females were significantly larger than males as expected, and no genotype effects were seen.

