## Supplementary figures and images for "Stochasticity in Dietary Restriction-Mediated Lifespan Outcomes in *Drosophila*"

### Supplementary Figure 1

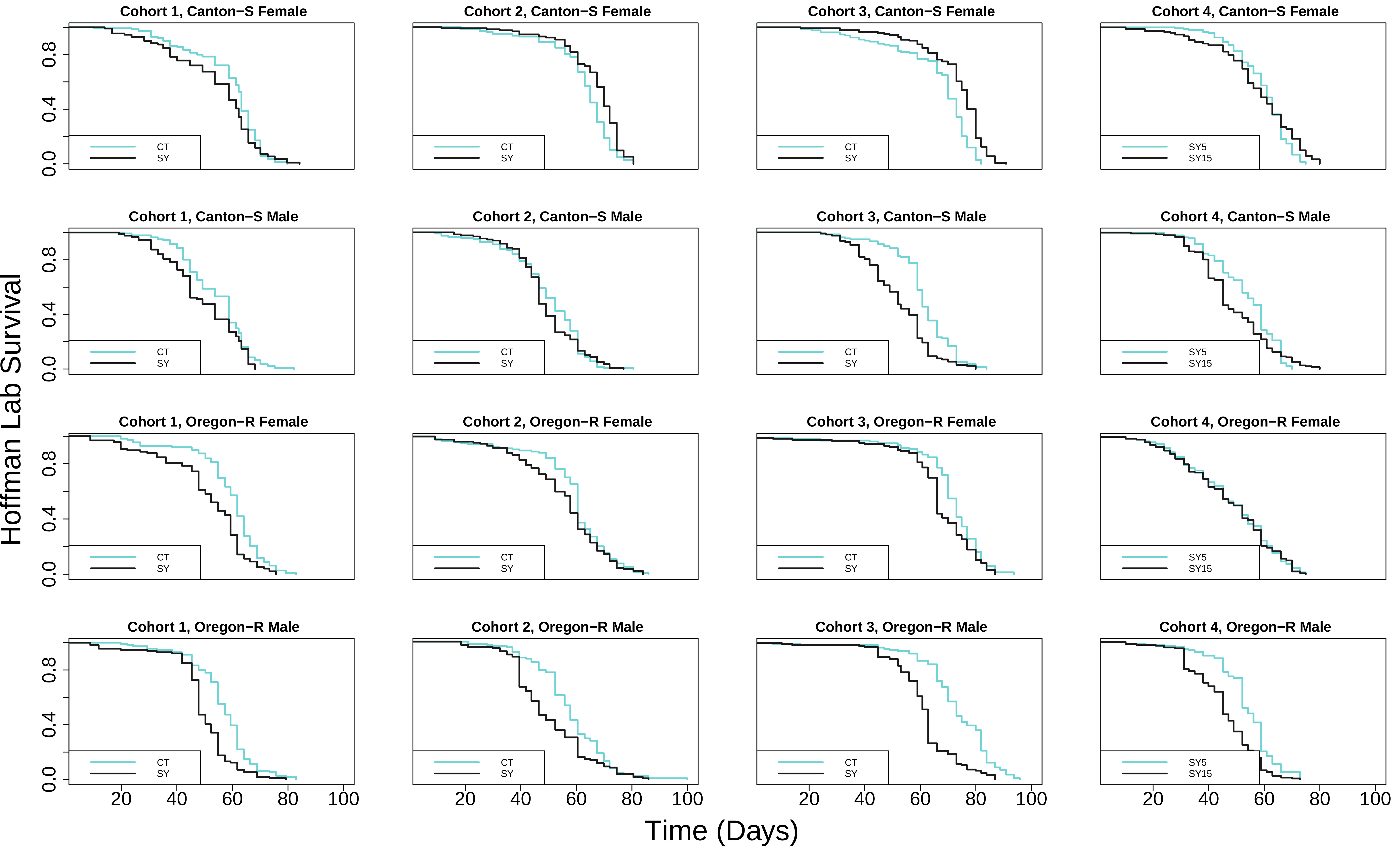

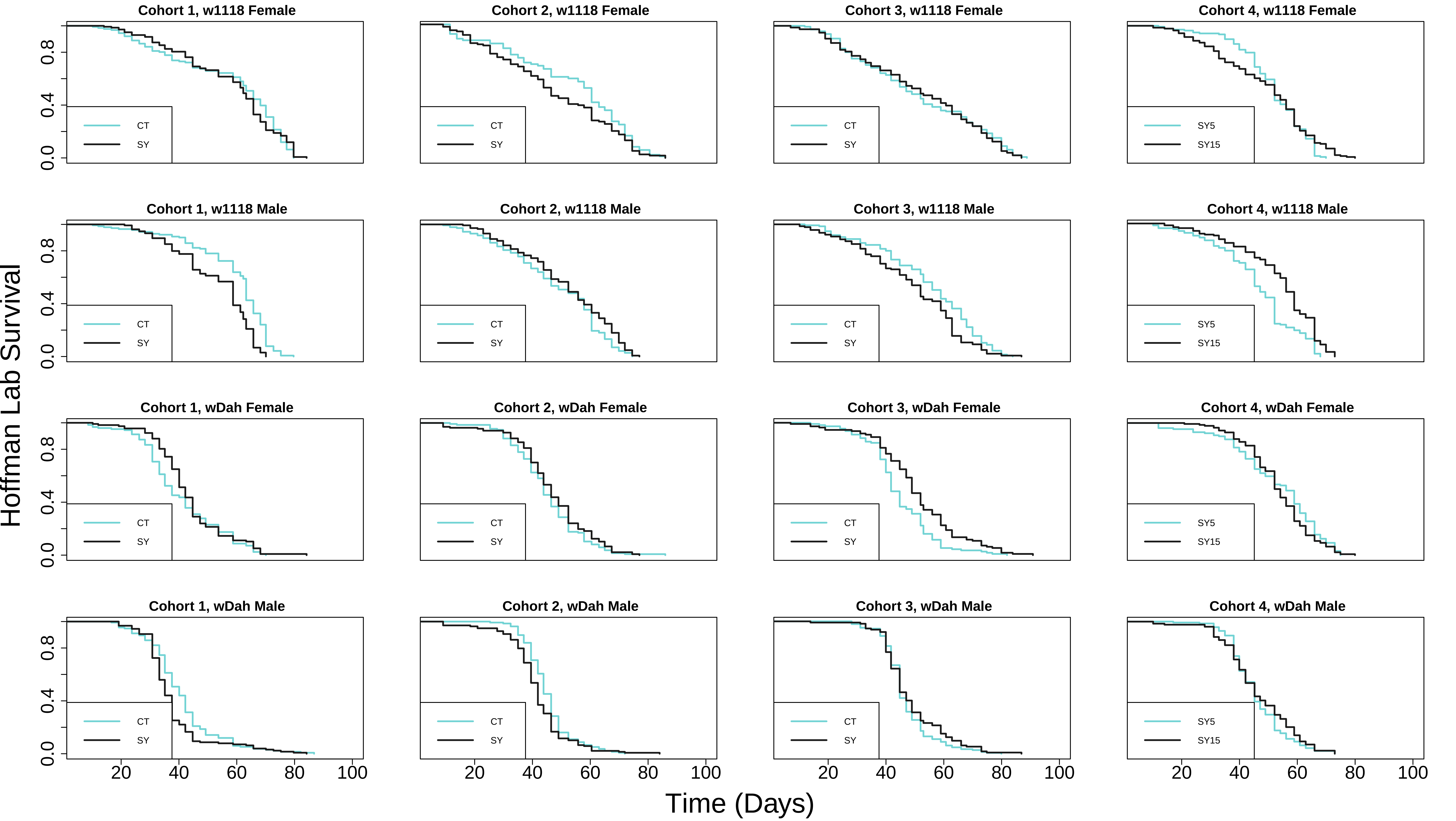

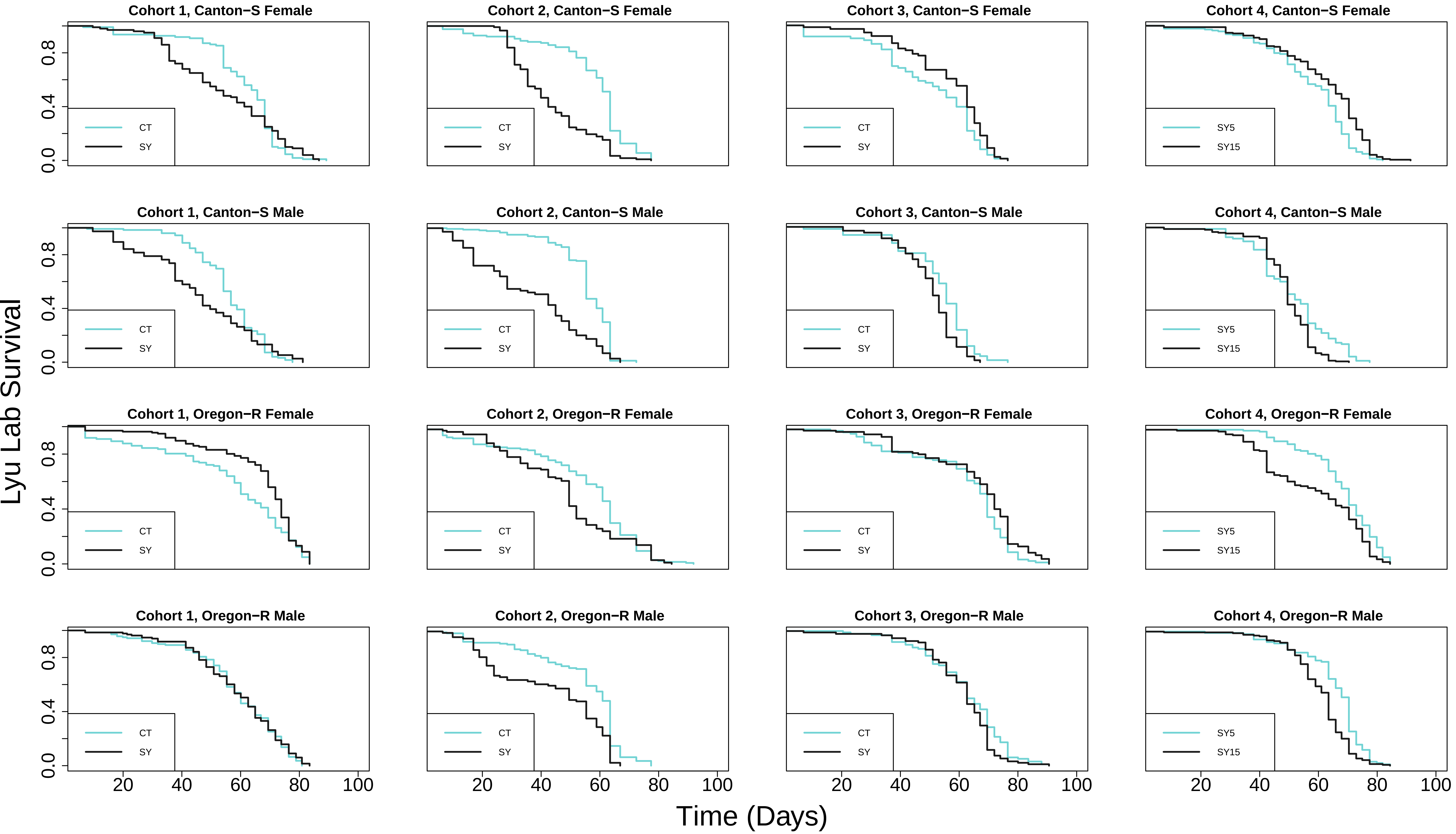

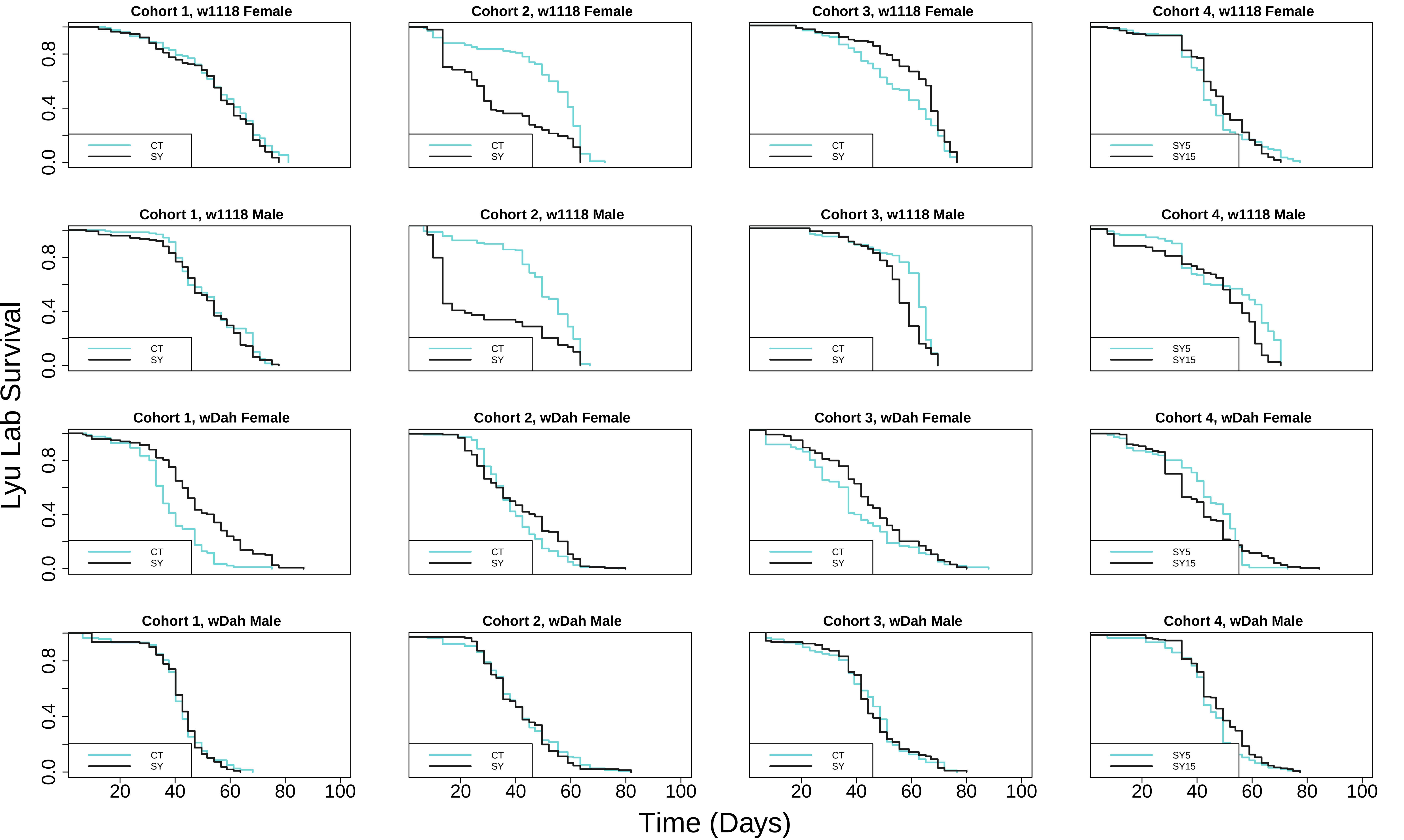

### Supplementary Figure 2

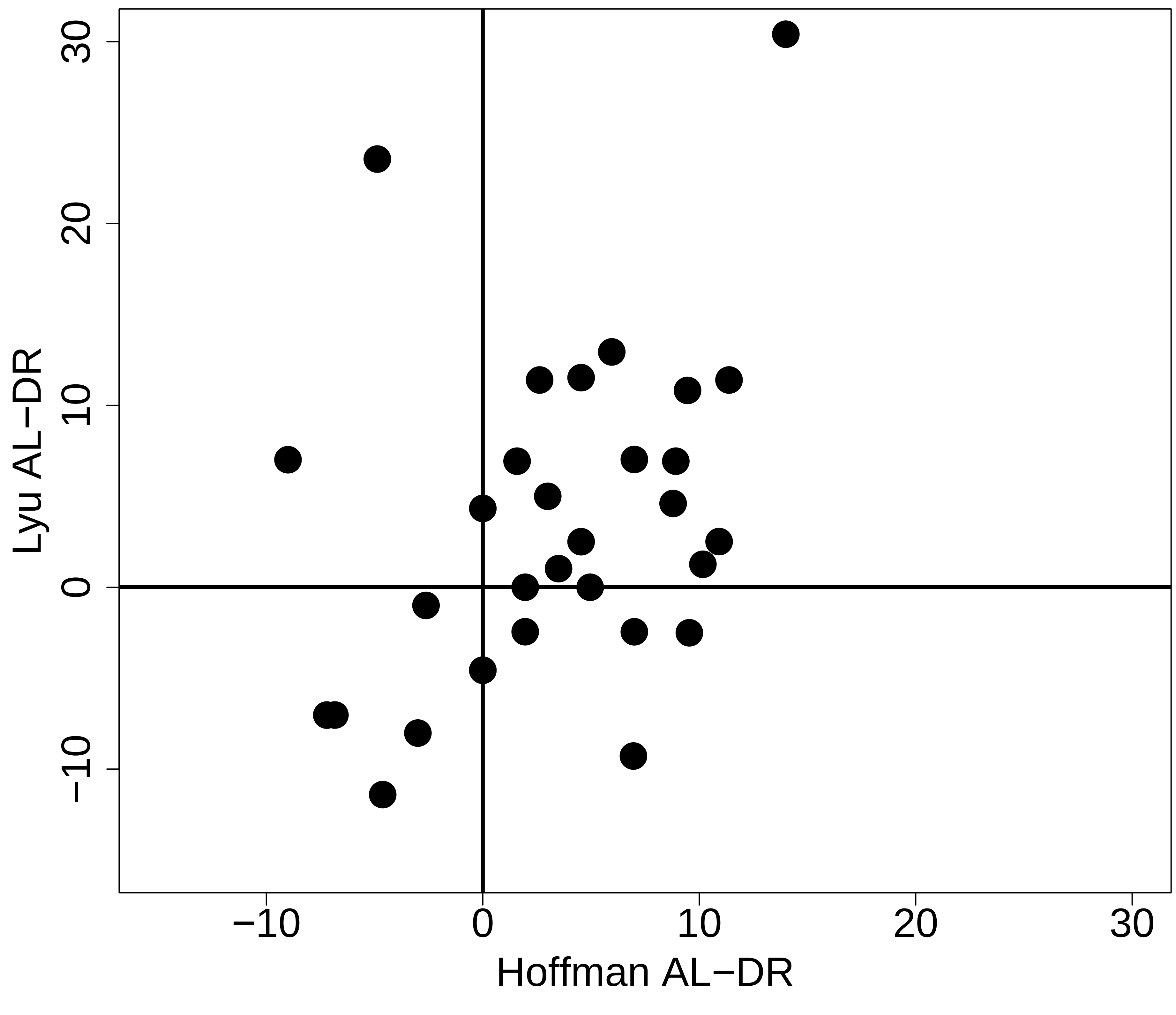

### Supplementary Figure 3

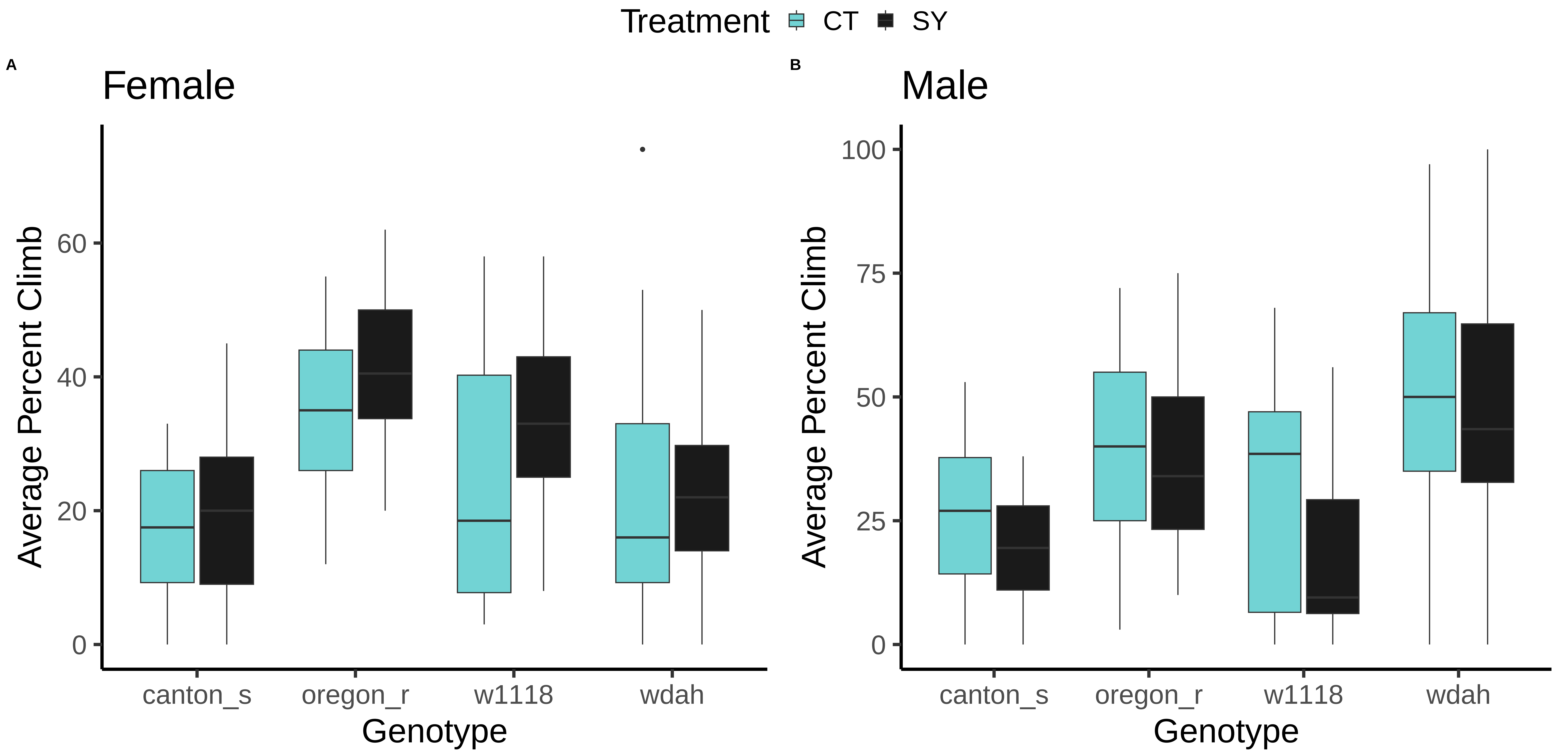

### Supplementary Figure 4

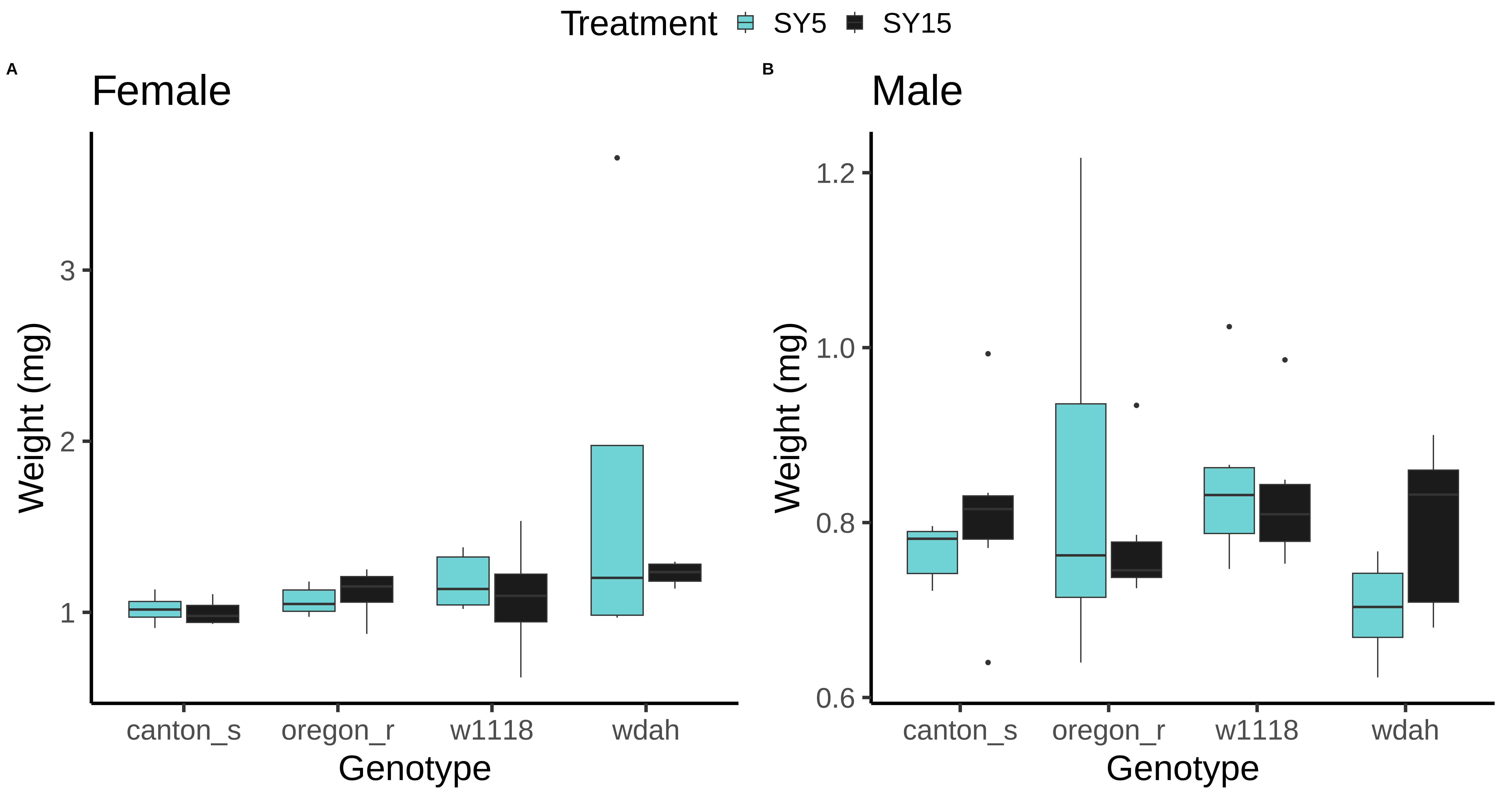
