## Supplementary Table 2 for "Stochasticity in Dietary Restriction-Mediated Lifespan Outcomes in *Drosophila*"

**Supplementary Table 2. Cox-Snell $R^2$ Values for full and reduced models to assess individual covariate contributions.**

| Factor | Cox-Snell $R^2$ (Cohort 1-4) |
|---|---|
| Full model | 0.1678643954 |
| Reduced model (-Lab) | 0.1673727175 |
| Reduced model (-Cohort) | 0.1434511787 |
| Reduced model (-Diet) | 0.1667438220 |
| Reduced model (-Sex) | 0.1461134875 |
| Reduced model (-Genotype) | 0.0615816753 |
